# CALM-VLM: CALIBRATION AND SELECTIVE PREDICTION IN VISION–LANGUAGE MODELS FOR RELIABLE BRAIN MRI CLASSIFICATION

**DOI:** 10.64898/2026.04.10.717865

**Authors:** Nikhil J. Dhinagar, Chirag Jagad, Pavithra Senthilkumar, Sophia I. Thomopoulos, Mahir H. Khan, Sook-Lei Liew, the ENIGMA-Stroke Recovery Working Group, Nerisa Banaj, Michael R. Boric, Lara A. Boyd, Amy Brodtmann, Jessica M. Cassidy, Adriana B. Conforto, Steven C. Cramer, Adrienne N. Dula, Fatemeh Geranmayeh, Chris M. Gregory, Brenton Hordacre, Abhishek Jaywant, Steven A. Kautz, Kristan A. Leech, Martin Lotze, Maria Mataró, Fabrizio Piras, Emily R. Rosario, Nerses Sanossian, Heidi M. Schambra, Nicolas Schweighofer, Na Jin Seo, Surjo R. Soekadar, Gregory T. Thielman, Carolee Winstein, George F. Wittenberg, Kristin A. Wong, Paul M. Thompson

## Abstract

Recent advances in vision-language models (VLMs) have demonstrated strong multimodal capabilities for medical image analysis. However, their confidence in diagnostic predictions is often unclear, limiting adoption in clinical settings. We introduce CALM-VLM (**CAL**ibration **M**echanism for Vision-Language Models), which integrates confidence calibration and selective prediction into a generative 3D VLM. To create CALM-VLM, we fine-tuned the Med3DVLM architecture for Alzheimer’s disease (AD) and stroke classification as initial test cases. To improve reliability, we incorporated temperature scaling on the VLM’s generative outputs. The calibrated model then selectively abstained from predictions when uncertain; this also improved its diagnostic accuracy. Experiments across multi-site MRI datasets, from 10 countries worldwide, show that CALM-VLM improved confidence relative to uncalibrated VLMs. Coverage-adjusted test receiver-operator characteristic curve-area under the curve (ROC-AUC) increased by 5% to 13% for both diagnostic tasks across independent test sets. Our calibrated VLM achieved a test ROC-AUC of 0.951 for AD classification and 0.905 for stroke classification. These findings highlight the importance of calibrated, uncertainty-aware VLMs for trustworthy neuroimaging AI.

## 1. Introduction

VLMs trained on medical imaging and text data have emerged as powerful frameworks for unified reasoning across data modalities, achieving state-of-the-art results in image/text retrieval, medical visual question answering (VQA), captioning, classification, and even generating reports [35] [36]. Despite these advances, their *uncertainty calibration* - the alignment between predictive confidence and correctness - remains an under-explored frontier, particularly for decoder-only architectures that output free-form text. In clinical and research contexts such as diagnostic radiology, overconfident classifications could lead to inaccurate diagnoses. While confidence calibration methods such as temperature scaling [1] [2] and conformal prediction [3] have improved reliability in stand-alone discriminative models, such as convolutional neural networks (CNNs) and vision transformers (ViTs), they have not yet been extended to generative VLMs, where predictions are output in the free-form text sequences (conditional probability for next token prediction) rather than discrete class probabilities.

### Related Work

#### Vision Models for brain image analysis

Deep convolutional networks (CNNs) [4] and vision transformers (ViTs) [5] have achieved strong performance across neuroimaging tasks, including the detection of Alzheimer’s disease [4] and Parkinson’s disease [6], based on brain MRI. Hybrid architectures such as the Swin Transformer (SwinT) [6] and DeComposed Former (DCFormer) [7], have handled 3D image inputs while maintaining efficiency.

#### Medical Vision-Language models

VLMs are powerful multimodal frameworks that can relate medical images to associated text. In medical imaging, extensions of Contrastive Language-Image Pretraining (CLIP) [33], Bootstrapping Language-Image Pretraining (BLIP) [34], Large Language and Vision Assistant (LLaVA) [21] have been adapted for radiology [8], [9], [10]. Recent medical AI agents [11] [12] build upon the reasoning capabilities of Large Language Models (LLMs) to assist with complex clinical tasks [13]. Despite their versatility, existing medical VLMs rarely address uncertainty calibration or the reliability of their text outputs [14], features that are critical to clinical applications.

#### Calibration of AI models

Model calibration aims to align predicted confidence with the true likelihood of being correct. Early calibration techniques, such as parametric Platt scaling and non-parametric isotonic regression, were originally developed for traditional classifiers [15].

Temperature scaling [2] is a simple yet effective post-hoc calibration technique for deep probabilistic models. More recent work has applied conformal prediction [16] and selective prediction [17], and extended calibration to multimodal settings [3].

In this work, we apply VLMs to 3D volumetric T1-weighted brain MRIs to classify Alzheimer’s disease and stroke - two common brain conditions where reliability and calibrated model confidence are critical for AI-assisted radiologic diagnosis. VLMs can be utilized in different capacities - retrieval, classification, report generation [12]. Here we utilized it akin to a classifier, to learn a joint image/language feature space, and benefit from the constraining nature of the language component of the VLM as opposed to noisy image only features in vision encoders.

Although a greater range of diagnoses are seen in practice, we start with two common conditions (Alzheimer’s disease and Stroke) to study model performance in a controlled setting. We propose CALM-VLM, a calibrated 3D VLM that selectively predicts only when it can confidently make a prediction. Our contributions include:

1. Fine-tuned multimodal VLM for neuroimaging through domain-specific prompts.
2. Introducing temperature scaling-based confidence calibration to align confidence with correctness.
3. Implementing a selective prediction mechanism that allows the model to provide diagnostic answers to questions only when confident.
4. Benchmarking against vision-only baselines for Alzheimer’s disease and stroke classification.

This work bridges the gap between predictive accuracy and model confidence, offering one of the first systematic studies of calibration and selective prediction in VLMs for clinical neuroimaging applications.

## 2. Methods

### 2.1 Model Architecture

#### Vision-only Baseline

We trained the classic 3D DenseNet121 CNN [18], a well-known workhorse for neuroimaging AI applications. We also trained a DCFormer [7] - a 3D convolutional transformer-hybrid architecture designed to capture both local and global context in volumetric medical images. The DCFormer provides a computationally efficient middle ground between CNNs and ViTs [5].

#### Vision-Language Model (VLM)

We finetuned Med3D-VLM [19], which combines a 3D DCFormer [7] with the Qwen-2.5-7B Language Model [20]. The architecture follows the LLaVA [21] paradigm:

#### A. Contrastive Vision Encoder

1. The 3D vision encoder (DCFormer) [7] converts the T1-w MRI volume (128×256×256) into a sequence of visual tokens.

#### B. Projector

The multimodal projector facilitates interaction between visual and text tokens. The projector adopts the MLP-mixer architecture, where MLP is Multi-layer Perceptron [22]. In parallel, the input text is tokenized using the vocabulary of the LLM. The resulting image and text tokens are then fused and passed to the LLM for joint reasoning.

#### C. Transformer Decoder

We used the cutting-edge Qwen2.5-7B-Instruct [20], pretrained for instruction-following tasks. The LLM is fine-tuned using Low-rank adaptation (LoRA) [23] to significantly reduce the number of trainable parameters.

The model is trained using a VQA format. Each training example consists of:

1. Image: T1-weighted MRI scan
2. Question: “*What is the diagnosis for this {age}-year-old {sex} based on the MRI scan?*”. We created several variants of this template to provide natural augmentation to the text prompts to improve generalization.
3. Answer: single-token ground truth (“AD” or “stroke” or “control”).

During inference, the VLM outputs a probability distribution over the entire vocabulary. To map these token-level probabilities to disease-level class probabilities (e.g., P(AD), P(stroke) and P(control)), we extract the logits of these diagnostic tokens. This converts the generative language model’s open-ended output space into a classification space suitable for calibration.

### 2.2 Training and Adaptation Strategies

#### A. Vision encoder pretraining

We used the default Med3DVLM weights for the 3D DCFormer – it had been pretrained on large-scale 3D CT images and paired text prompts using SigLIP. We also custom finetuned the vision encoder on our 3D MRI datasets (ADNI and ENIGMA-Stroke). We report the best-performing pretrained variants.

#### B. Efficient Fine-tuning

We varied the LoRA Rank and we finetuned the vision encoder and the projector. This reduced the trainable parameter count from 7 billion to 75 million.

### 2.3 Model Uncertainty

#### 2.3.1 Calibration error metrics

##### A. Expected Calibration Error (ECE)

ECE [2] partitions prediction probabilities into *M* equally-spaced bins and takes a weighted average of the bins’ accuracy/confidence difference.

##### B. Brier Score [24]

This is calculated as the weighted sum of the squared differences between the predicted class probabilities and the one-hot encoded class labels over the entire test set.

#### 2.3.2. Calibration

We used *temperature scaling*, as it fits within the scope of this work, is fast and has inherently less risk of overfitting given it tunes a scalar ‘temperature’ parameter T>0 [2], which “softens” the softmax with T>1. T is optimized with respect to negative log-likelihood on the validation set. At test-time calibration is applied to the logits corresponding to the diagnostic tokens of interest (AD, stroke, and healthy control). This better aligns the confidence of the model based on the prior alignment with respect to the correctness on the validation set.

#### 2.3.3 Selective Prediction: We evaluated selective prediction - where the model abstains when confidence falls below a given threshold τ, for τ ϵ [0.5, 0.95]. We reported accuracy-confidence reliability curves and identified optimal thresholds for target accuracy levels for fixed coverage (i.e., prediction rate)

## 3. Experimental Setup

### 3.1 Datasets

We used the publicly available ADNI (Alzheimer’s Disease Neuroimaging Initiative) [25] and OASIS (Open Access Series of Imaging Studies) [26] brain MRI datasets for AD classification. In this work, the dementia diagnosis in ADNI and OASIS3 is referred to as AD, following the clinical definition of AD rather than the more recent research framework defining biological AD by amyloid positivity (Aβ+) [27]. ADNI was split into 2,577, 302, and 1,219 T1-w scans for training, validation, and testing, respectively, with unique subjects restricted to a specific fold to avoid data leakage given the longitudinal data. 600 scans from OASIS were used for zero-shot testing. For stroke classification, we used data collected by the Enhancing Neuroimaging Genetics through Meta-Analysis (ENIGMA) Stroke Recovery working group [28] in 10 countries worldwide. The ENIGMA-Stroke dataset is one of the largest multisite retrospective stroke datasets to date [28]. For stroke classification, we used 1077, 120, and 300 T1-w scans for training, validation, and testing, respectively. We used 83 held-out scans from a single site for zero-shot testing. **Table 1** summarizes the datasets used in this work. In the current paper, we do not include experiments on multiclass classification, as the available training data were imbalanced with respect to age. This design choice is consistent with recent AI studies that primarily focused on disease-specific fine-tuning [29] [30].

**Table 1.** Summary of data used in this work for training and testing. *one subject’s sex in the stroke dataset was unavailable. Datasets used include ADNI (Alzheimer’s Disease Neuroimaging Initiative), OASIS (Open Access Series of Imaging Studies) for Alzheimer’s disease classification and data collected by the Enhancing Neuroimaging Genetics through Meta-Analysis (ENIGMA) Stroke Recovery working group for stroke classification.

| Dataset | N scans<br>(N subjects) | Age range<br>(years) | N Sex<br>(F/M) | N Sites |
| --- | --- | --- | --- | --- |
| ADNI | 4,098<br>(1,188) | 55.2-95.8 | 556F/<br>632M | 62 U.S. sites |
| OASIS | 600<br>(600) | 43.5-97.0 | 341F/<br>259M | 1 U.S. site |
| ENIGMA<br>Stroke | 1,580<br>(1,580) | 17.0-93.0 | 616F/<br>963M<br>* | 50 sites from the U.S.,<br>Italy, Canada, Brazil,<br>Norway, UK,<br>Germany, Spain,<br>Australia, and China |

### 3.2 Data Pre-processing

In line with similar studies [4], all 3D T1-weighted brain MRI scans were pre-processed via standard steps for neuroimaging analyses including: ‘skull-stripping’, N4 bias field correction [31], linear registration to a template [32] with 9 degrees of freedom, and isotropic resampling of voxels to 2-mm resolution. All images were *z*-transformed. T1-w scans were re-sized to 128×256×256 voxels before model training to match the default training set-up for Med3DVLM.

### 3.3 Training

We conducted a random search to select hyperparameters for vision-only and the VLM models. Models were all trained for 5 epochs. Training was performed using DeepSpeed ZeRO-2 to enable memory-efficient distributed optimization.

### 3.4 Evaluation

We evaluated our models with ROC-AUC, and accuracy for the two classification tasks. We also used three standard machine translation metrics: (1) bilingual evaluation understudy (BLEU); (2) Recall-Oriented Understudy for Gisting Evaluation - Longest Common Subsequence (ROUGE-L), to evaluate overlap between generated result and the ground truth, and (3) BERTScore to quantify the similarity between the two based on semantic embeddings.

## 4. Experimental Results

**Tables 2 and 3** compare the performance of vision-only models with our proposed CALM-VLM for AD and Stroke. **Table 4** presents calibration results for the VLM under different LoRA ranks, including error metrics (ECE, Brier Score). Calibration and selective prediction significantly improved model reliability. **Figure 2** shows how calibration aligns model confidence with its true correctness - the calibrated (*green*) curve is spread across the full range.

**Table 2.** Performance of the CALM-VLM Across Datasets (ADNI and independent site OASIS) for Alzheimer’s Disease Classification. The top model results are represented using bold-face formatting.

| Model | Calibrated | Coverage | ADNI ROC-AUC □ | ADNI ROUGE □ | ADNI BLEU □ | ADNI BERT-Score □ | OASIS 0-shot ROC-AUC □ |
| --- | --- | --- | --- | --- | --- | --- | --- |
| DenseNet 121 CNN | - | 1.00 | 0.834 | - | - | - | 0.849 |
| DCFormer | - | 1.00 | 0.904 | - | - | - | 0.778 |
| VLM-75M | - | 1.00 | 0.932 | 0.847 | 0.847 | 0.998 | 0.848 |
| **CALM-VLM-75M** | **✓** | 0.88 (0.80 for 0-shot) | **0.951** | - | - | - | **0.889** |

**Table 3.**
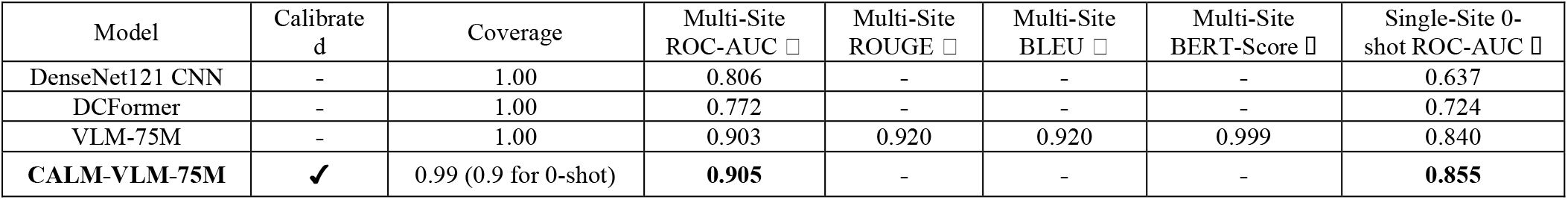
Performance of the CALM-VLM Across the ENIGMA-Stroke Dataset (multi-site and independent single-site) for Stroke Classification.

| Model | Calibrated | Coverage | Multi-Site ROC-AUC □ | Multi-Site ROUGE □ | Multi-Site BLEU □ | Multi-Site BERT-Score □ | Single-Site 0-shot ROC-AUC □ |
| --- | --- | --- | --- | --- | --- | --- | --- |
| DenseNet121 CNN | - | 1.00 | 0.806 | - | - | - | 0.637 |
| DCFormer | - | 1.00 | 0.772 | - | - | - | 0.724 |
| VLM-75M | - | 1.00 | 0.903 | 0.920 | 0.920 | 0.999 | 0.840 |
| **CALM-VLM-75M** | **✓** | 0.99 (0.9 for 0-shot) | **0.905** | - | - | - | **0.855** |

**Table 4.** Comparative Analysis of Calibration Error and Low-Rank Adaptation Settings.

| Model | Coverage | LoRA Rank | ADNI ROC-AUC □ | ADNI Calibration Error (ECE) □ | ADNI Brier Score □ | OASIS 0-shot ROC-AUC □ |
| --- | --- | --- | --- | --- | --- | --- |
| VLM-75M | 1.00 | 4 | 0.932 | 14.51% | 0.145 | 0.848 |
| **CALM-VLM-75M** | 0.88 (0.80 for OASIS) | 4 | **0.951** | 6.12% | **0.112** | 0.889 |
| VLM-75M | 1.00 | 8 | 0.895 | 13.6% | 0.139 | 0.879 |
| CALM-VLM-75M | 0.18 (0.23 for OASIS) | 8 | 0.926 | **4.16%** | 0.116 | **0.966** |

**Figure 1.**
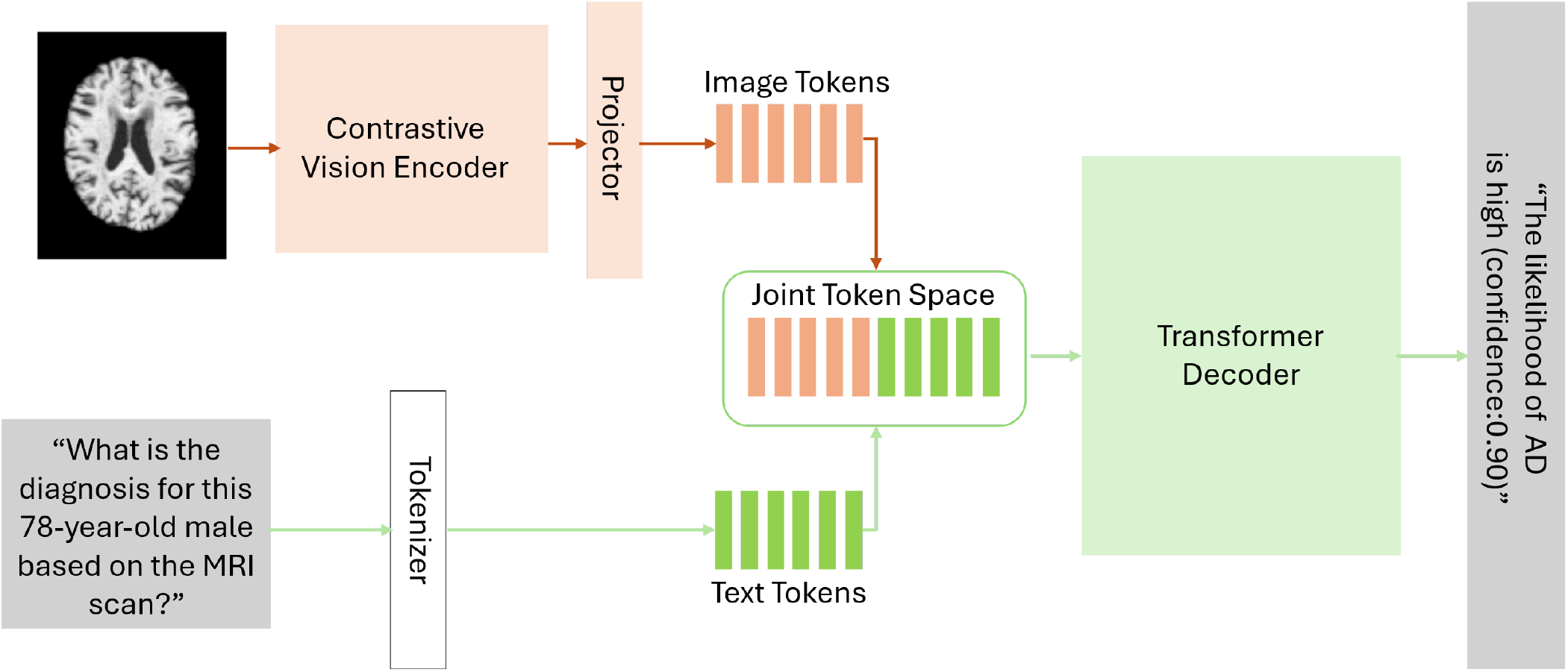
CALM-VLM, our 3D medical Vision-Language Model. Input: T1-w MRI and the text prompt: “*What is the diagnosis for this 78-year-old male based on the MRI scan?*”. Output: ‘*The likelihood of Alzheimer’s disease is high (confidence: 0*.*90)*”. The model reports the predicted diagnosis and its calibrated confidence score - communicating uncertainty in this open-ended VQA setting.

**Figure 2.**
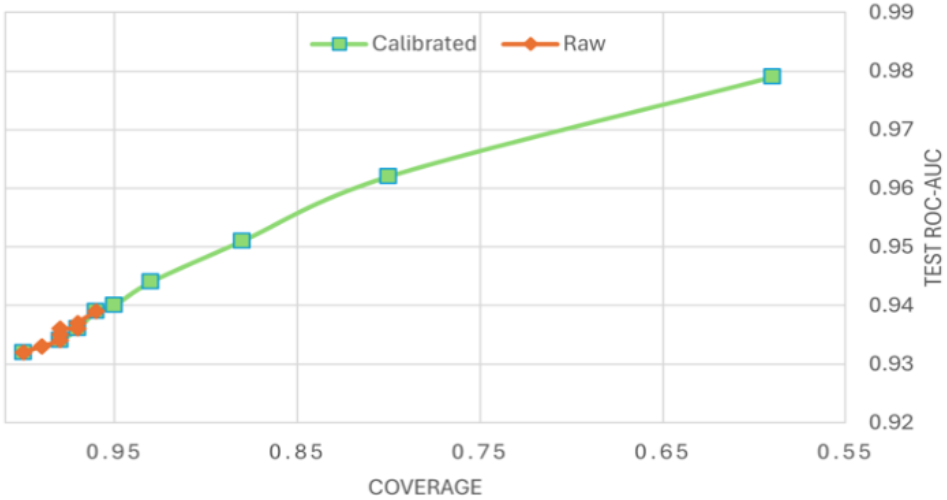
Test ROC-AUC vs. coverage for calibrated (*green*) and raw (*orange*) VLM models. Calibration spreads confidence scores across the full range, enabling higher test-time performance by abstaining on low-confidence predictions (selective prediction). Coverage is the prediction rate of the model.

## 5. DISCUSSION

### 5.1 Key findings

#### A. VLM improved classification accuracy

The calibrated vision-language model achieved 5% and 10% higher ROC-AUC compared to the 3D vision-only baselines for AD and stroke classification. This suggests that integrating subject-specific demographic context (age, sex) through natural language helps models make more informed predictions. The VLM in our experiments typically outperformed on the 0-shot out-of-distribution dataset/site compared to the vision-only models (5% for AD and 13% for stroke for ROC-AUC

#### B. Calibration improves model confidence

After calibration, the VLM’s confidence was better aligned with its predictive accuracy. The calibrated CALM-VLM achieved ROC-AUC of 0.951 for AD classification with a coverage of 0.88 - meaning that it can reliably auto-diagnose over 88% of the unseen test subjects with high accuracy, while the remaining cases can be reviewed by an expert.

#### C. Patient context capability

Demographic factors are important confounders for AD. The VLM learned to weigh these context factors appropriately, whereas vision-only models cannot access this information.

### 5.2 Clinical Implications

#### A. Trustworthy AI-assisted diagnosis

Overconfidence due to poor model calibration is a major barrier to AI adoption in clinical applications. Our results show that VLMs can be trained to make accurate predictions along with trustworthy confidence estimates. This is a key development and is critical for clinician trust and patient safety. As an example, a neuroradiologist can focus on uncertain cases based on model confidence.

### B. Explainability through language

VLMs natively support the use of demographic context via natural language to complement medical images, improving interpretability.

#### 5.3 Limitations and Future Work

Future extensions will test a wider range of conditions including other calibration methods to assess the generalizability and adaptability of the proposed CALM-VLM framework. In the current work, for purposes of simplicity, we focused on two binary classification tasks (Alzheimer’s disease versus healthy, and stroke versus healthy), as these are common benchmarks where accuracy is routinely reported in medical image analyses; this makes it easier to tell how a method’s accuracy relates to other similar methods. However, in future we will study a more realistic scenario where people are assigned to the most likely diagnosis among 3 (or more) categories, using techniques to assess N-way classifiers. A slight variant of this approach is to include individuals who have multiple diagnoses (e.g., both Alzheimer’s and stroke), and permit outputs to report all likely diagnoses and their confidence. In the current training data, it cannot be ruled out that at least some of the patients had either Alzheimer’s disease or mild cognitive impairment prior to the stroke. A more detailed evaluation would separate these cases in the training and test data, and evaluate the model’s accuracy specifically in these cases.

For subjects with multiple possible diagnoses, our framework could be adapted to identify the co-occurring conditions, and quantify uncertainty across alternatives. The reason we did not address this case in the current report is that our Alzheimer’s and stroke datasets are somewhat imbalanced for age, which could lead to “short-cut learning” where the classifier uses the patient’s age and sex as part of the diagnostic decision, as it is associated with diagnosis in the training data. Future work will address this issue, as the patient’s age and sex should affect the class-conditional priors for disease classification, but the methods to incorporate this should still be tested on data that are matched for age and sex, to avoid short-cut learning.

A further clinical observation is that the ENIGMA-Stroke dataset is mainly assessing chronic stroke, which is easier to identify on T1-weighted MRI than acute stroke. Future work that includes acute stroke patients in the training and test data will be needed to assess the range of accuracy of the method as a function of time since stroke. With enough training data, it should be possible to stratify the evaluation by time since stroke, and by lesion volume and location, which are all likely to influence accuracy; other training data, such as atrophy, microbleeds, prior infarcts, etc., may further affect accuracy.

In the future, we will also incorporate additional neuroimaging modalities, biomarkers, genotype, clinical, and electronic health record data. Additional modalities are especially important for the diagnosis of acute stroke and for distinguishing it from other common neurological conditions. Our proposed CALM-VLM system could be interfaced with radiological information systems, to integrate model outputs and confidence scores into clinical workflows.

## 6. Conclusion

We presented CALM-VLM, a calibrated, uncertainty-aware VLM for reliable brain MRI diagnosis. Our approach combined temperature scaling and confidence-based selective prediction. CALM-VLM improved confidence calibration and accuracy compared to the vision-only baselines. This work is a step towards trustworthy foundation models in neuroimaging, that can both answer diagnostic questions and know when to answer.

## 7. Acknowledgments

This work was supported by the U.S. National Institutes of Health, under NIA grant U01 AG068057 andS10OD032285. We are grateful to the ENIGMA Stroke Working Group (https://enigma.ini.usc.edu/ongoing/enigma-stroke-recovery/) for providing the stroke dataset.

